# Advancement in Cellular Topographic and Nanoparticle Capture Imaging by High Resolution Microscopy Incorporating a Freeze-Drying and Gaseous Nitrogen-based Approach

**DOI:** 10.1101/2023.09.28.559906

**Authors:** Kunihiro Uryu, Nadine Soplop, Timothy P. Sheahan, Maria-Teresa Catanese, Chuong Huynh, John Pena, Nancy Boudreau, Irina Matei, Candia Kenific, Ayako Hashimoto, Ayuko Hoshino, Charles M. Rice, David Lyden

## Abstract

Scanning electron microscopy (SEM) offers an unparalleled view of the membrane topography of mammalian cells by using a conventional osmium (OsO_4_) and ethanol-based tissue preparation. However, conventional SEM methods limit optimal resolution due to ethanol and lipid interactions and interfere with visualization of fluorescent reporter proteins. Therefore, SEM correlative light and electron microscopy (CLEM) has been hindered by the adverse effects of ethanol and OsO_4_ on retention of fluorescence signals. To overcome this technological gap in achieving high-resolution SEM and retain fluorescent reporter signals, we developed a freeze-drying method with gaseous nitrogen (FDGN). We demonstrate that FDGN preserves cyto-architecture to allow visualization of detailed membrane topography while retaining fluorescent signals and that FDGN processing can be used in conjunction with a variety of high-resolution imaging systems to enable collection and validation of unique, high-quality data from these approaches. In particular, we show that FDGN coupled with high resolution microscopy provided detailed insight into viral or tumor-derived extracellular vesicle (TEV)-host cell interactions and may aid in designing new approaches to intervene during viral infection or to harness TEVs as therapeutic agents.

## Introduction

Scanning electron microscopy (SEM) has enabled the viewing of fine ultrastructures associated with external cellular boundaries. However, water-rich (∼80% by volume) cells surrounded by a lipid-rich membrane present a challenge in achieving optimal preservation of topography at high resolution due to the requirement for dehydration and maintenance of membrane fatty acids without compromising membrane properties. Earlier SEM protocols using ethanol resulted in suboptimal preservation of membranes (Mitchell, 1969; Ongun et al., 1968). This led to exploration of alternative methods, including the application of osmium tetra-oxide (OsO_4_) (Saunders et al., 1968) and of derivatives of other organic solvents (Saunders et al., 1968), as well as lowering the temperature of reagents (Dallam, 1957). To date, the combination of OsO_4_ post-fixation and ethanol dehydration has been widely accepted and the adverse effects of this method on membrane integrity are considered relatively minor (Dallam, 1957; Saunders et al., 1968).

The desire to visualize the relationship between functional molecules and ultrastructures spurred the development of correlative light and electron microscopy (CLEM). CLEM has been widely deployed transmission electron microscopy (TEM) (Avinoam et al., 2015; Bykov et al., 2016; de Boer et al., 2015; Gupta et al., 2019; Hampton et al., 2017; Peddie et al., 2014). In contrast, SEM CLEM for topographical analysis has been hindered by the adverse effects of ethanol and OsO_4_ on retention of fluorescence signals. To overcome this problem, cryo-SEM or environmental SEM imaging was employed which eliminated the use of organic solvents or OsO_4_ to retain fluorescent signals. However, either approach prevents direct interactions between the electron beam and the sample surface due to the presence of ice, extra-thick conductive coating, or moisture on the sample surface (Albornoz et al., 2020; Strnad et al., 2015). Furthermore, the low vacuum with the presence of gas in the SEM chamber limits resolution (Hempel, 2017; Joy and Joy, 1996). Alternative methods were introduced to utilize hydrated samples for SEM CLEM (Thiberge et al., 2004; Wojcik et al., 2015), and demonstrated a clear relationship between intracellular elements and fluorescent signals. These approaches have improved the ability to view intracellular structures as well as fluorescent signals of reporting molecules. However, precise imaging of cell surfaces to elucidate the relationship between pathogens or other extracellular particles and host cells in high resolution remains a challenge.

Here, we developed a new sample preparation method free of organic solvents or OsO_4_ to minimize the adverse effects on membrane topography and applied a freeze-drying method using gaseous nitrogen, FDGN, for SEM and SEM CLEM. We demonstrate that samples prepared by FDGN maintain robust fluorescent signals via fluorescent light microscopy (FLM) and exceedingly detailed topography via SEM. To test the capabilities of FDGN, we assessed hepatitis C virus (HCV)- and tumor-derived extracellular vesicles (TEVs)-human cell interactions *in vitro*. With FDGN, our SEM-based analyses suggested how membrane structures are organized and interact with HCV or TEV. We applied the FDGN method to other imaging applications, including a super-resolution scanning microscopy called helium ion microscopy (HIM) to visualize the cell surface without conductive coating at even higher resolution (Joens et al., 2013) and TEM CLEM to connect SEM CLEM view to intracellular views. Evaluation of the data generated from multi-imaging systems improved our understanding of the molecular and structural organization of the cells in qualitative and quantitative manners. Incorporating FDGN processing provided an opportunity to develop mechanistic insight into how viral pathogens and TEVs influence the structure of the host cell membrane to allow entry.

## Results

### Detailed cell membrane architecture post HCV infection of Huh7.5 cells

We first illustrated FDGN workflow (Fig. 1) and below describe the examination of the structural morphology of a model system of HCV- or TEV-host cell interaction using various methods to evaluate FDGN technology.

**Fig. 1.**
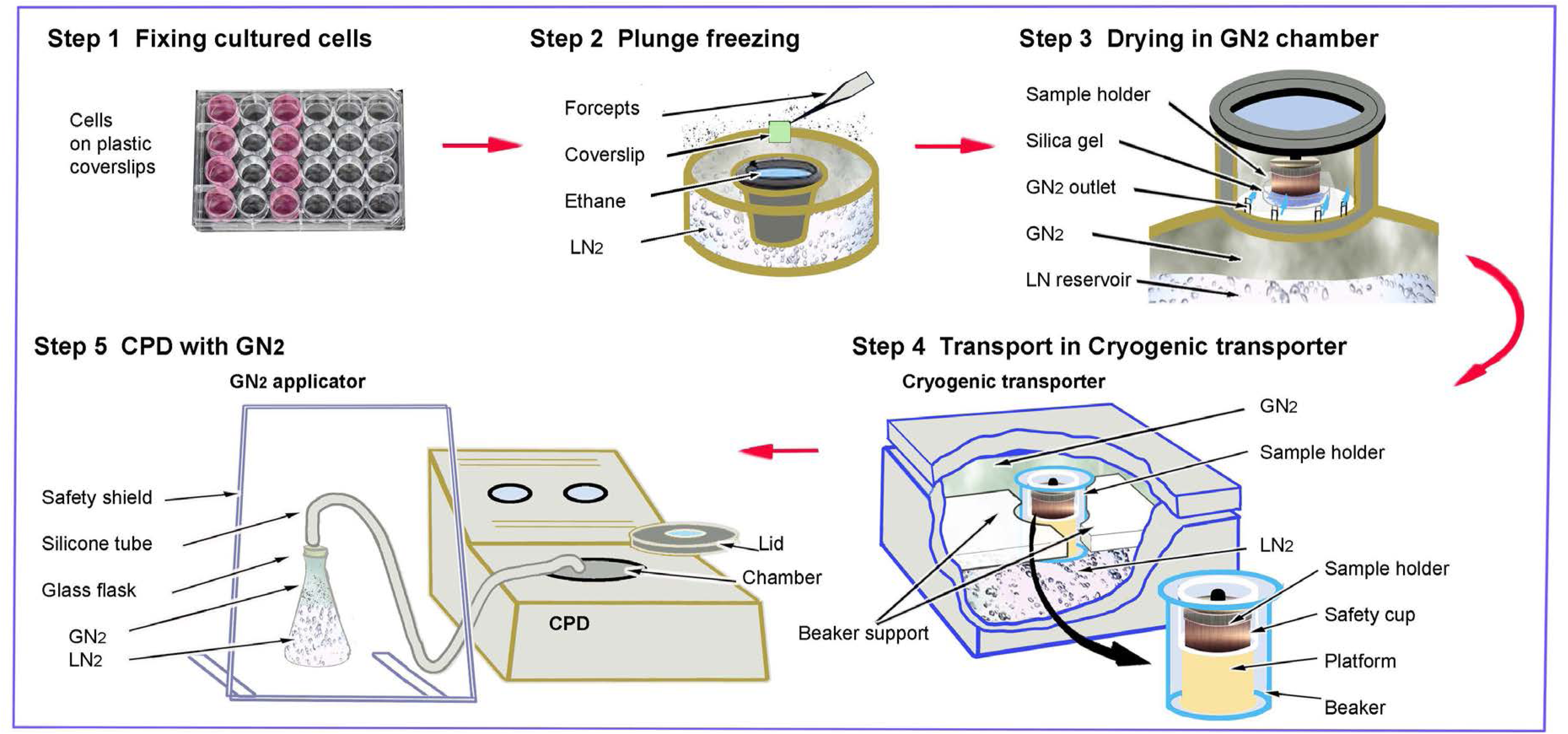
Workflow of FDGN. **Step 1. Fixing cultured cells.** Cells attached to gridded plastic coverslips were treated in culture wells according to the study’s design and fixed with an EM fixative. **Step 2. Plunge freezing.** An approximately 3 x 3 mm-sized *coverslip* was held by a pair of *forceps*, and the liquid covering the coverslip was carefully withdrawn by filter paper. The coverslip was frozen by plunging into liquid *ethane* pre-cooled by *LN_2_*. **Step 3. Drying in GN*_2_* chamber.** The coverslip was placed in a pre-cooled sample holder in a drying chamber, which was connected to an LN_2_ reservoir and supplied *GN_2_* to the chamber through the outlet to keep the chamber moisture free and at an aimed temperature between −120 and −100 C° overnight. The presence of a Petri dish-filled *silica* gel offered additional support for the drying samples. **Step 4. Transport in Cryogenic tranporter.** A sample transport from the GN_2_ drying chamber to the *CPD* unit was done safely using a cryogenic tranporter made of a Styrofoam box. (The illustration shows the tranporter with a part of structure was removed to reveal the inside.) In the tranporter, a glass beaker containing a metal platform was placed, and its position was firmly fixed by two arms of the beaker support made of Styrofoam. The bottom half of the unit was filled with *LN_2_* and the upper half was filled by *GN_2_*. The sample holder was placed in a plastic safety cup, transferred from the drying chamber to the transporter, located on a metal platform, and outfitted with a lid. In this way, the sample was surrounded by cold *GN_2_* during the transportation but avoided direct contact with *LN_2_.* **Step 5. CPD with GN_2_.** For the successful *CPD* procedure, a *GN_2_* applicator was arranged. The glass flask filled with *LN_2_* half way and gently placing a topple connecting to a silicone tube made flexible application of cold GN_2_ flow available. Using this applicator, the CPD chamber was pre-cooled by introducing GN_2_. The sample holder was translocated to the CPD chamber and a ceaseless flow of *GN_2_* was applied by holding the free end of the silicon tube immediately over it. A flow of GN_2_ was continuously applied to the sample holder in the chamber during the process of the lid closure, and upon completion, the programmed CPD operation was initiated. A safety shield was used as a personal protection equipment. Note: Throughout the procedure, it was critical for the areas of tools and devices to contact the sample holder to be pre-cooled by LN_2_ or GN_2_ from the *GN_2_* applicator. (See Materials and Method for more details.)

Human-derived hepatoma Huh-7.5 cells at 24-hour post-infection (24 hpi) were prepared using one of the following four methods: 1) conventional SEM with OsO4 post-fixation (CSEM), 2) FDGN with critical point drying (FDGN), 3) conventional SEM method without OsO_4_ post-fixation (CSEM-Os), and 4) FDGN without critical point drying (CPD) as a control (CPD-ctr) (summarized in Table 1). SEM at a high magnification can demonstrate the characteristic features of the samples prepared from each method (Fig. 2a).

**Fig. 2.**
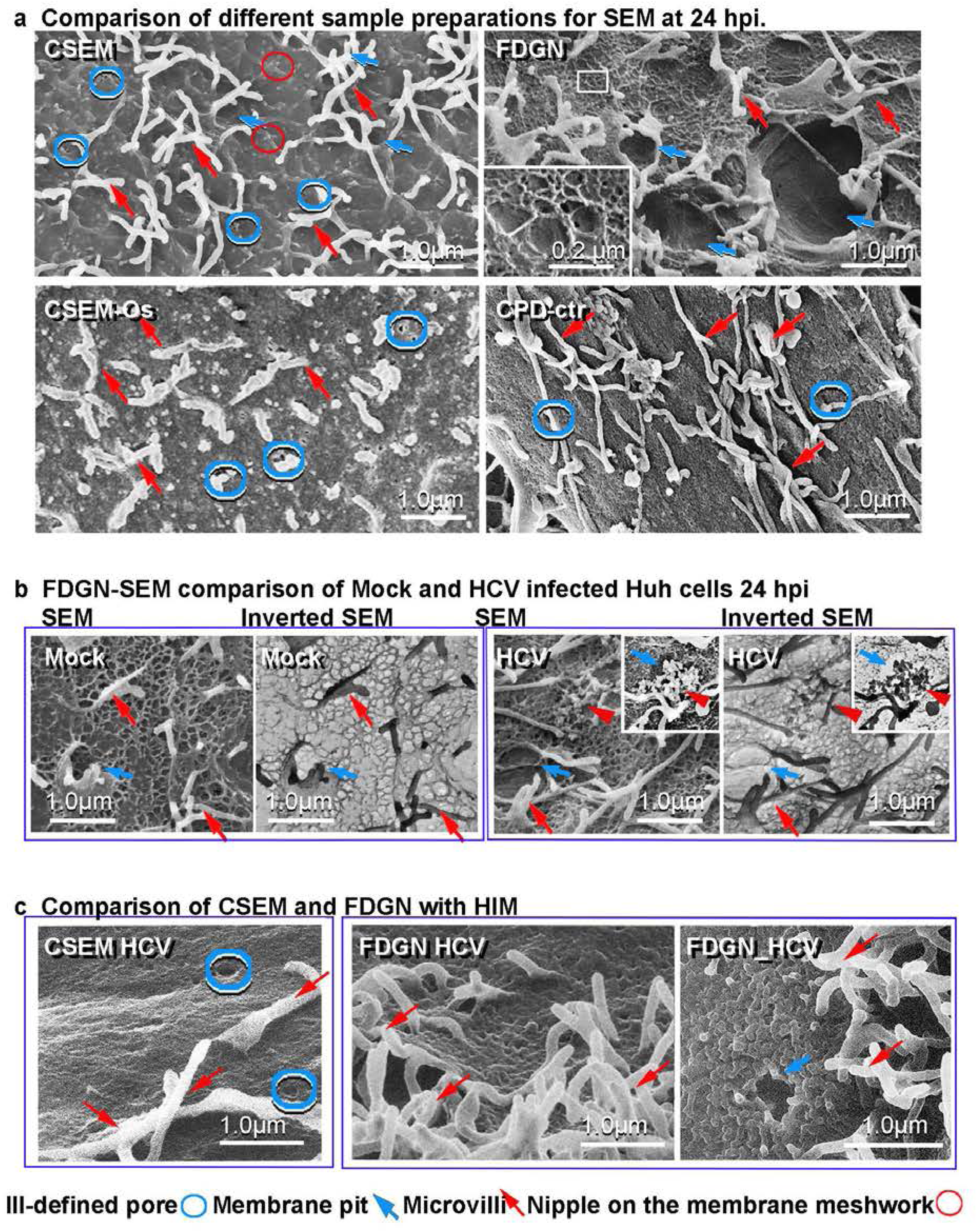
FDGN provides superior preservation of cellular architecture in Huh7.5 cells. **a,** Comparison of different sample preparations for SEM at 24 hpi. CSEM image (***upper left***) reveals fine profiles of microvilli (***red arrow*** 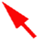) and smooth membranes with irregular, ill-defined pores (***blue circle*** 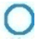). Fine stipples and associated nipples were highlighted (***circle*** 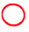). FDGN image (***upper right***) shows microvilli (***red arrow*** 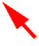), extensive meshwork, and membrane pits (***blue arrow*** 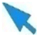). CSEM-Os image (***lower left***) reveals deformation and poor preservation of the membrane. CPD-ctr image (***lower right***) reveals aggregated microvilli and a distinct textured membrane. **b,** FDGN-SEM comparison of mock and HCV infected Huh7.5 cells at 24 hpi. SEM and Inverted SEM images of mock cell (***left panel***) reveal short microvilli, membrane pits (***blue arrow*** 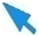), and a relaxed meshwork and circumventing membrane pits without distortion. At 24 hours post HCV infection, Huh7.5 cells (***right panel***) display elongated microvilli, increased number of membrane pits (***blue arrow*** 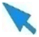) and a reorganized meshwork circumventing the membrane pits. The presence of HCV particles (***arrowhead*** 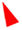) and its position within the membrane pits are indicated (***inset***). **c,** Comparison of CSEM and FDGN by HIM. CSEM image shows a featureless cell surface with microvilli (***red arrow*** 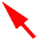), and ill-defined pores (***blue circle*** 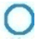) (***left panel***). FDGN processed sample shows improved preservation of a detailed membrane meshwork with microvilli (***red arrow*** 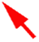) (***middle panel***). Higher magnification image shows the elaborate meshwork organization at a membrane pit (***blue arrow*** 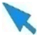) (***right panel***).

**Table 1.** Methods tested for microscopy analyses of mammalian cells.

| Method tested | Method abbreviation | Key treatments | Microscopy system tested |  |  |  | Pros | Cons |
| --- | --- | --- | --- | --- | --- | --- | --- | --- |
|  |  |  | FLM | SEM | HIM | TEM |  |  |
| Conventional SEM including CPD | CSEM | ethanol, osmium tetroxide, CPD | - | ++ | ++ | ++ | Osmium mitigates deformation of topography caused by ethanol. | No fluorescence is visible and questionable structure preservation for high resolution. |
| Conventional SEM without osmium tetroxide | CSEM-Os | ethanol, CPD | - | - | - | + | Provides good dehydration. | No fluorescence is visible, and membrane is poorly preserved. |
| Freeze Dry with Gaseous Nitrogen with CPD | FDGN | nitrogen gas, CPD | +++ | +++ | +++ | +++ | Retains fluorescent signals and lipid membrane structure. | Controlled operation without expensive equipment. |
| CPD omitting control for Freeze Dry with Gaseous Nitrogen | CPD-ctr | nitrogen gas | +++ | + | + | n/a | It retains fluorescent signals. | Imperfect dehydration. |

CSEM revealed well-defined views of the membrane topography decorated with microvilli (*red arrow*) (Fig. 2a*, CSEM, upper left panel*). The membrane exhibited a smooth texture with fine fibrous stipples (<20nm in thickness) which were connected (*red circle*). We observed membrane pits as cup shaped membrane depressions with reduced appearance of a cytoskeleton meshwork, but the cups (*blue arrow*) were often not well exposed in this sample. There were a number of minute ill-defined pores in a wide range of a sizes (mostly >30 nm in diameter) that appeared randomly distributed throughout the cell (*blue circle*). Next, we evaluated the FDGN method, which showed remarkably fine details of the cell topography, including microvilli (*red arrow*), a meshwork consisting of an extensive number of fine fibers, also connected with fine nipples (<30nm in diameter) (*red circle, highlighted in the inset*). and overt membrane pits (*blue arrow*)(Fig. 2a, *FDGN, upper right panel*). The clear presentation of the fibrous meshwork and pits were unique to the samples prepared by FDGN as compared to standard CSEM or other methods employed here. CSEM-Os revealed thick and short microvilli (*red arrow*) on the rough surface along with irregularly spaced minute pores (*blue circle*) (Fig. 2a, *CSEM-Os, lower left panel*). No meshwork, including nipples, nor membrane pits were observed, suggesting deformation and damage to the membrane integrity. When CPD was omitted from FDGN, cells exhibited fine microvilli (*red arrow*) that were unanimously flattened and aggregated, thus failing to reveal how they extend into the external environment (Fig. 2a*, CPD-ctr, lower right panel*). The cell membrane displayed a detailed surface that regularly consisted of numerous pores (*blue circle*). However, this sample lacked the stipples, fibrous meshwork, or membrane pits that were clearly observed with FDGN. As the CPD-ctr omitted application of liquid carbon dioxide to remove water, it is likely that this corrupted membrane presentation was due to imperfect dehydration caused by surface water (Nordestgaard and Rostgaard, 1985).

Comparing the samples prepared by four different dehydration methods, we concluded that FDGN offered the finest detail of cell surface elements. FDGN extensively preserved a fine meshwork with fibers and nipples, possibly representing the sub-membrane cytoskeletal network and associated protein complexes, respectively, and the fine demonstration of close linkage between membrane pits and HCV virus or TEV with the cell, which will be described further below. In contrast this meshwork and membrane pits were less well preserved and fewer details could be observed using CSEM, CSEM-Os, or CPD-ctr methods.

### Effects of Huh7.5 cell Meshwork by HCV infection

HCV infection is known to cause phenotypic changes with a dynamic rearrangement of the cell membrane following infection by TEM (Blanchard and Roingeard, 2018; Egger et al., 2002). Since SEM studies were limited to non-HCV infection (Caldas et al., 2020; Cortese et al., 2017), we carfully examined the cell topography, meshwork, and membrane pits using FDGN SEM in either mock-infected or HCV-infected cells at 24 hpi. Mock-infected cells displayed short and sporadic microvilli (Fig. 2b, *Mock, SEM, left panel, red arrow*), well-organized meshwork and membrane pits (*blue arrow*). A monochromatic inversion of the SEM micrograph (Fig. 2b, *Inverted SEM*) highlighted the relationship between the meshwork and membrane pits (*blue arrow*) and revealed the regularity of the meshwork organization.

The FDGN SEM images of HCV-infected cells revealed reorganized cell surface with an increased number of microvilli (*red arrow*) and membrane pits (*blue arrow*) (Fig. 2b, *24 hpi, SEM, right panel*). Elongated microvilli could also be seen. The clusters of particles (between 50 and 70 nm in diameter) represent the HCV virions tethered to the cell surface (*red arrowhead*). A cluster of particles appeared in the proximity of a membrane pit (*inset*). Monochromatic inversion of the SEM (Fig. 2b, *24 hpi, Inverted SEM, right panel*) highlighted the distorted arrangement of the meshwork, which appeared to be compressed due to newly formed membrane pits. A membrane pit (*blue arrow*) surrounded the cluster of particles (*red arrowhead*) (*inset*). These images, obtained using FDGN, revealed a rearrangement of the cell topography following HCV virus-host cell interactions which included reorganized meshwork, elongated microvilli, increased number of microvilli and increased number of membrane pits.

### Application of FDGN to HIM

Next, we compared CSEM or FDGN using super-resolution HIM. Although CSEM provided a clear view of microvilli (*red arrow*) and surface of the cells, CSEM also revealed ill-defined pores (*blue circle*) and a smooth texture on the membrane surface (Fig. 2c, *CSEM*, *left panel*). In contrast, FDGN preserved the fine microvilli (*red arrow*) and meshwork of stipples covering the cell surface (Fig. 2c, *FDGN, middle panel*). The details of these stipples were strikingly highlighted at a type of membrane pit (*blue arrow*) (Fig. 2c, *FDGN, right panel*). The FDGN HIM images confirmed the observations made using SEM, in that CSEM compromised cell topography while FDGN retained superior detail.

### Nuclear re-arrangment post HCV infection

Huh7.5 cells were transfected with a red fluorescent protein-nuclear localization sequence-mitochondrially-tethered interferon-beta promoter stimulator protein 1 (RFP-NLS-IPS) construct (*RFP*) (Jones et al., 2010), thus HCV infection of the cell triggers nuclear translocation of RFP. We compared HCV- or mock-infected Huh7.5 cells at 24 hpi using FDGN and fluorescent light microscopy (FLM). FDGN preserved both fluorescent signals of the nuclear marker (*NclBlue*) and RFP remarkably well in Huh-7.5 cells at 24 hpi. Mock cells indicated persistent distribution of RFP in mitochondria (Fig 2a*, FLM*, Mock), while a subset (37.54 ± 12.17%) of the infected cells displayed nuclear translocation (*arrow*) (Fig 2a*, FLM, HCV*), and the remaining infected cells showed the RFP signal in mitochondria. We subsequently tested FDGN for SEM CLEM. First, we identified nuclear RFP (*arrow*) in a subset of HCV-infected cells using FLM (Fig 2a, *SEM CLEM, FLM, right panel*). This region of interest (ROI) was examined with SEM, and thereafter FLM and SEM images were merged (Fig 2a, *SEM CLEM, merged, right panel*). One of the two recognized cells displaying RFP in this field (*inset*) is shown at a higher magnification (Fig 2a, *SEM CLEM, Close-up, right panel*) and revealed a bulged nuclear (*Nucl*) profile. This bulged nuclear phenotype was observed in approximately 30% of the cells with nuclear RFP.

This nuclear phenotype was further examined. Mock-infected Huh7.5 cells presented flat cell profiles with unrecognizable nuclei from the external view (Fig 2b*, Mock, Upper left*). Following HCV infection, bulged nuclei, with horizontal or global expansion of the cell nuclei, were observed. Nuclear blebs at the periphery of the nucleus and evidence of nucleus compartmentation were often observed with horizontal expansion (Fig 2b*, HCV 24 hpi, upper middle panel*). A type of bulged nuclei, observed following the global expansion, displayed a variety of forms, including numerous membrane pits (*blue arrow*) (Fig 2b*, HCV 24 hpi, upper right panel*), wavy or spiky membranes with numerous microvilli (Fig 2b*, HCV 24 hpi, lower left panel*), or ruptured nuclear envelopes and cell membranes (Fig 2b*, HCV 24 hpi, lower middle panel*). Ruptured nuclear envelopes occasionally appeared in extended nucleoplasm (*Bleb*), where nuclear pores (*circle*) (*inset*) could also be seen (Fig 2b*, HCV 24 hpi, lower right panel*). These phenotypes suggest a dynamic rearrangement of nuclear organization following HCV infection. Application of FDGN in conjunction with SEM CLEM with RFP-NLS-IPS labelled cells revealed a specific phenotype of nuclear bulge, which occurred along with simultaneous structural changes, including nuclear reorganization and cell membrane rearrangements.

### Quantitative analyses of microvilli and membrane pits

High-quality images obtained using FDGN allowed us to apply semi-quantitative data analyses of cell morphology in response to HCV infection *in vitro*. First, we evaluated the density of microvilli under different conditions, including mock infection and HCV infection with bulged nuclei (HCV+BN) and without bulged nuclei (HCV-BN) (Fig. 3c*, Microvilli Density, far left panel*). Compared to mock-infected cells, microvilli density increased in HCV+BN cells but not in HCV-BN cells (mock: 8.26 ± 2.50 /µm^2^; HCV-BN: 7.19 ± 1.39 /µm^2^; HCV+BN: 17.37 ± 1.47 /µm^2^; p-value < 0.05 for Mock vs. HCV+BN as well as HCV-BN vs. HCV+BN).

**Fig. 3.**
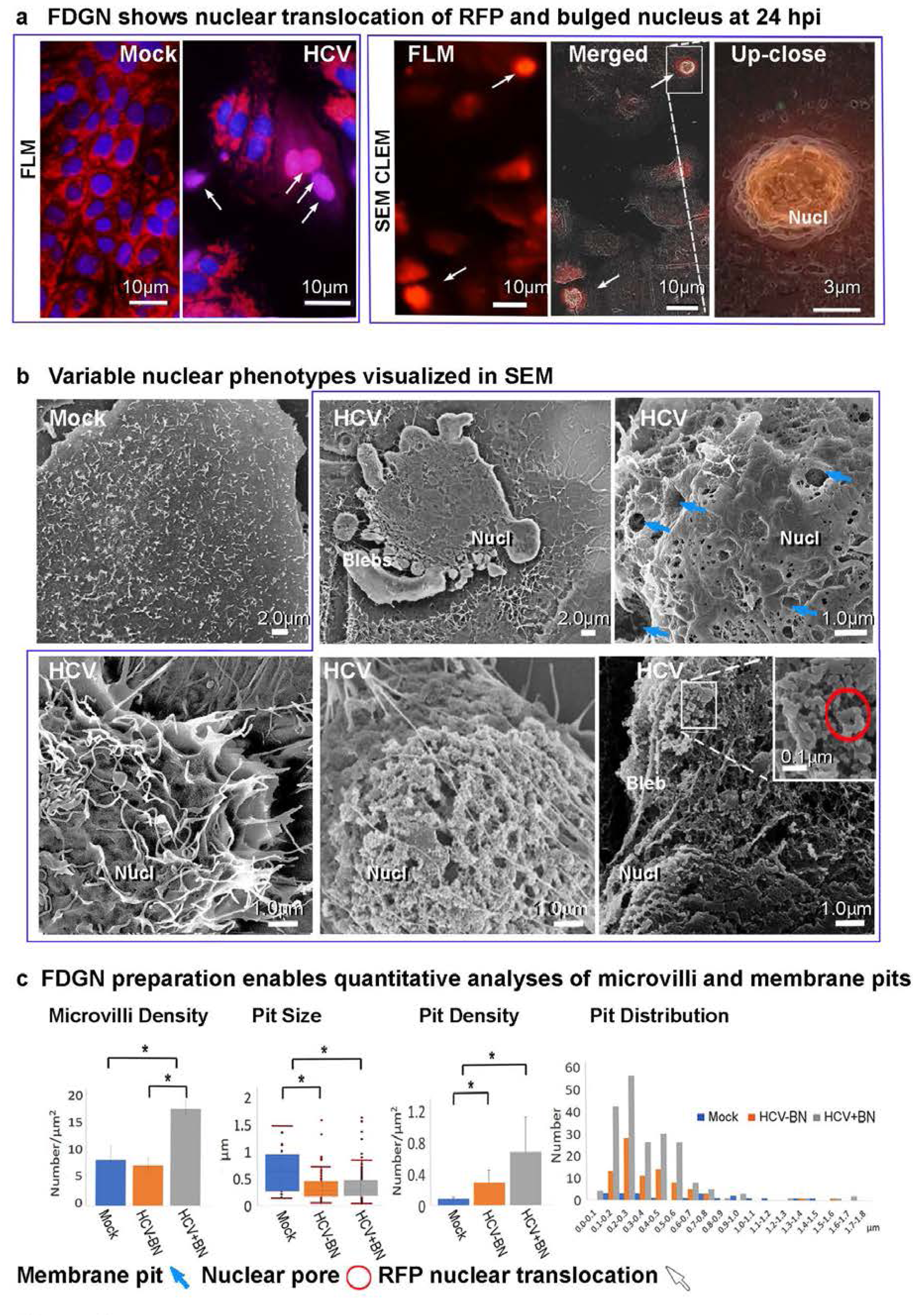
The impact of HCV infection on the nuclei of Huh7.5 cells revealed using FDGN. **a, FDGN** reveals nuclear translocation of RFP and altered nuclear phenotype at 24 hpi **FLM** (***left panel***) shows retention of both **NucBlue** and **RFP** fluorescent signals at 24 hpi. In the **Merged** (***left panel***) view, a group of cells in the center show translocation of RFP to the nucleus (***arrow ↑***), and another subset of cells at the upper right show the mitochondria-associated RFP. In **SEM CLEM**, **FLM** (***right panel***) shows RFP positive cells in a field. The combination of FLM and SEM views of the ROI were shown in **Merged** (***right panel***) view, and the **arrows** (***↑***) show two representative cells with nuclear RFP. Higher magnification of a cell (***inset***) reveals bulged and blebbing nuclear (***Nucl***) profiles (***Up-close, right panel***). **b,** Variable nuclear phenotypes visualized with SEM A low power view of **Mock** infected Huh7.5 cell shows a typical flat cell profile; and the nucleus is not prominent (***upper left panel***). A blebbing nucleus with horizontal expansion (***upper middle panel***), global nuclear expansion with numerous membrane pits (***blue arrow*** 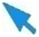) (***upper right***), or wavy membrane with numerous microvilli (***lower left***) were all observed. The nuclear expansion can be associated with a major exposure of the nucleoplasm (***lower middle panel***) or a partial exposure of the nucleoplasm with its blebbing expansion (***lower right panel***). A higher magnification view (***inset***) shows nuclear pores (***circle*** 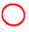). Scale bars were added accordingly. **c,** FDGN preparation enables quantitative analyses of membrane pits A quantitative comparison of the **Microvilli Density** (per µm^2^), **Pit Size** (diameter by µm) and **Pit Density** (per µm^2^) of membrane pits between the mock infected or HCV infected Huh cells with or without bulging nuclei. Mock, HCV-BN, and HCV+BN groups are shown. The size distribution of membrane pits is also presented. * Statistically significant, with p-value < 0.05.

Next, we compared membrane pits in the same three groups of Huh7.5 cells. Membrane pits in mock-infected cells were larger than those in HCV-BN or HCV+BN cells (Fig. 3c*, Pit Size, 2^nd^ left panel*) (Mock: 0.66 ± 0.39 µm; HCV-BN: 0.39 ± 0.25 µm; HCV+BN: 0.38 ± 0.26 µm; p< 0.05 *for* Mock vs. HCV-BN and Mock vs. HCV+BN). Meanwhile, the density of membrane pits (per µm^2^), were lowest in mock-infected cells and increased in both HCV-BN and HCV+BN cells (Fig. 3c*, Pit Density, 2^nd^ right panel*) (Mock: 0.073 ± 0.006 /µm^2^; HCV-BN: 0.29 ± 0.16 /µm^2^; HCV+BN: 0.67 ± 0.44 /µm^2^; p-value < 0.05 for Mock vs. HCV-BN, and Mock vs. HCV+BN). Thus, following HCV-cell interactions, the size of membrane pits was reduced while the density of membrane pits on the surface increased, particularly in infected cells displaying bulged nuclei (Fig. 3c*, Distribution, far right panel*). These analyses confirmed a dynamic rearrangement of microvilli and cell membrane organization associated with HCV /cell interaction and its sequelae.

### Cell membrane architecture in WI-38 fibroblasts interacting with tumor-derived EVs

We also applied FDGN to investigate changes in human lung fibroblasts (WI-38 cells), as this cell line takes up TEVs shed by breast cancer cells and subsequent TEV uptake is linked to cancer metastasis (Hoshino et al., 2015). Initially, we compared WI-38 cells at 2 hours post TEV exposure (hpa) using the four methods described above for evaluation of Huh7.5 cellular topography using SEM at 10,000x magnification.

CSEM revealed fine microvilli (*red arrow*) and small (<200 nm; *small arrowhead*) and large TEVs (>200 nm; *large arrowhead)* (Fig. 4a*, CSEM, upper left panel*). The cell membrane displayed a unique, sheet-like texture. The cell surface exhibited an undulating profile with longitudinal hills, suggestive of the sub-membrane cytoplasmic cytoskeleton. The membrane also contained numerous ill-defined pores (>30nm; *blue circle)*. While a few membrane pits were noticeable, they were not fully exposed with this method (*blue arrow)*. Similar to Huh 7.5 cells, FDGN provided the most striking details of membrane topography in WI-38 fibroblasts. These included microvilli (*red arrow)*, small (*red small arrowhead*) and large TEVs (*red large arrowhead)*, and membrane pits (*blue arrow*) (Fig. 4a*, FDGN, upper right panel*). The cell surface was covered by a meshwork visible in exquisite detail that consisted of numerous fine fibers associated with minute nipples (>30nm) (*circle, highlited in the inset*). On the upper side of the image, the decorated fibers and nipples (>50nm) provided a view of the elongated meshwork (*dotted line*). Some of the larger TEVs displayed bumpy surfaces, which may represent clusters of small TEVs. In contrast, CSEM-Os resulted in a lack of detail and deformation of the cell surface, which was potentially masked by unfixed lipids. Different sizes of spherical structures, representing small (*small red arrowhead)* and large TEVs (*large red arrowhead)* were visible along with an ill-defined pore (*blue circle*) (Fig. 4a*, CSEM-Os, lower left panel*). Omitting CPD yielded observable microvilli (*red arrow)* as well as small (*small red arrowhead*) and large (*large red arrowhead*) TEVs (Fig. 4a*, CPD-ctr, lower right panel*). However, the microvilli were flattened and aggregated, and the meshwork seen with FDGN was not visible. Some poorly defined pores (*blue circle)* were also noticeable.

**Fig. 4.**
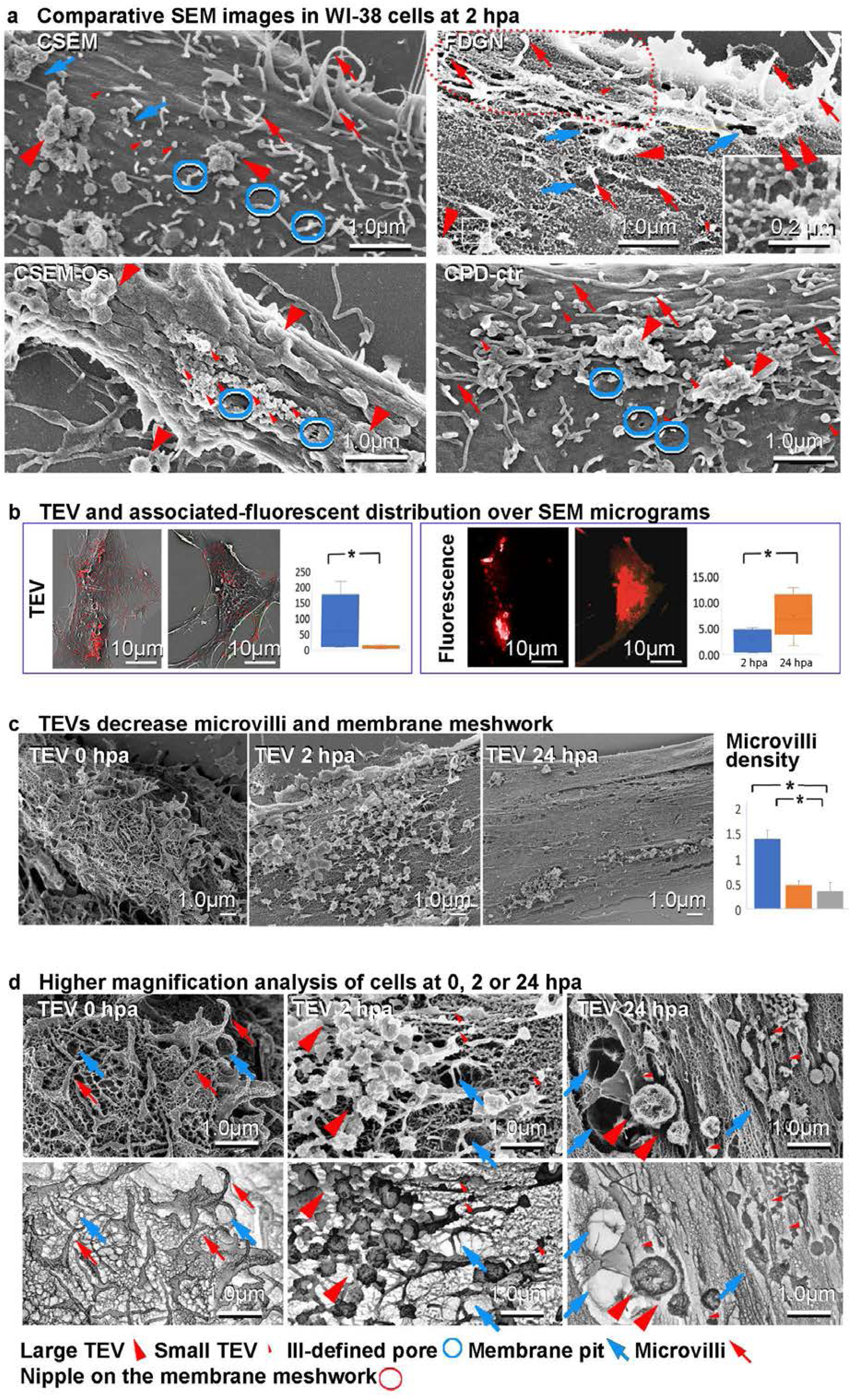
FDGN reveals detailed WI-38 cell organization. **a,** Comparative SEM images in WI-38 cells at 2 hpa with TEVs. CSEM (***top left***) revealed a detailed topography with microvilli (***red arrow*** 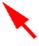), membrane pits (***blue arrow*** 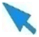), and ill-defined pores (***blue circle*** 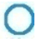) on the undulating membrane. Large TEV (***large arrowhead*** 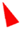) and small TEV (***small arrowhead*** 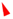) are visible in CSEM, and all other sample preparation methods. FDGN (***top right***) revealed a more detailed topography with microvilli (***red arrow*** 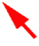), and membrane pits (***blue arrow*** 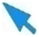). Fine fibers with associated nipples cover the cell surface (***circle*** 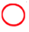). The dotted line highlights the meshwork comprised of fine fibers. CSEM-Os (***bottom left***) results in poor preservation and deformation of the membrane with and numerous pores (***blue circle*** 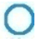). Omitting CPD (***bottom right***) results in flattened and aggregated microvilli (***red arrow*** 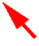) and numerous pores (***blue circle*** 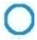). **b,** TEV and associated-fluorescent distribution in SEM micrographs. SEM micrograph showing distribution of FM1-43FX and PKH26 labled TEVs (***red dots*** 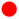) on the cell surface at **2 hpa** or **24 hpa (*left panel*)**. Histogram shows quantification of surface fluorescence at both **2 hpa** and **24 hpa**. Fluorescent microscopy of TEV associated fluorescence at **2 hpa** or **24 hpa** is shown (***right panel***). The histogram represents quantification of fluorescent areas by percentage. * Statistically significant with p-value < 0.05. **c**, TEVs decreased microvilli and mebrane meshwork WI-38 cells cultured with TEVs at **0 hpa** display numerous short microvilli and a relaxed meshwork (***left panel)***. At **2 hpa** cells exhibit a narrow more elongated shape with numerous TEVs tethered to the cell surface (***middle panel***). At **24 hpa** the narrow-elongated shape is maintained, and few TEVs remain on the cell surface and a tightly organized cytoskeleton can be observed (***right panel***). A quantitative comparison of the **Microvilli Density**(per µm^2^), at 0, 2, 24 hpa, is presented. **d,** Higher magnification analysis of cells at 0, 2 or 24 hpa **SEM** or inverted SEM views of WI-38 cells without TEV (**0 hpa**) or at **2** or **24 hpa** TEV exposure. Control WI-38 cells display an elaborate cell surface organization with numerous microvilli **(*red arrow*** 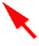), meshwork, and membrane pits (***blue arrow*** 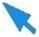) (***upper left panel***). The corresponding inverted SEM image highlights the relaxed arrangement of the meshwork and membrane pits (***blue arrow*** 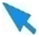) (***lower left panel***). At **2 hpa**, a SEM view reveals numerous small and large TEVs tethered to the cell surface via fibers (***upper middle panel***). The corresponding inverted SEM image reveals a reorganization of the membrane meshwork and membrane pits (***blue arrow*** 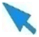) (***lower middle panel***). A number of large TEVs (***large arrowhead*** 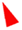) and small TEVs (***small arrowhead*** 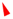) are tethered to the cell surface. SEM at **24 hpa** reveals a reduced number of small and large TEVs in the proximity of membrane pits (***upper right panel***). The cytoskeletal meshwork shows a directional orientation. The corresponding **Inverted SEM** image highlights a tight and directional meshwork, as well as TEV adjacent to a membrane pits (***lower right panel***).

In contrast to FDGN, the meshwork organization observed using CSEM also appeared to be masked by a layer of unfixed lipids covering the surface, along with numerous clusters of small pores, representative of corrupted meshwork fibers. CSEM images showed that only the large fibers of the meshwork were visible, whereas FDGN revealed long thick fibers (>30nm) as well as short thin fibers (<20nm in diameter) that ran in a parallel direction. As noted in our analyses with Huh7.5 cells, it appeared that the meshwork and nipples represented submembrane cytoskeleton and protein complexes, respectively, that undergo dynamic rearrangement as the TEVs intiate interaction with WI-38 cells. We therefore investigated TEVs and their interaction with WI-38 cells in more detail.

### TEV and fluorescent reporter distribution using SEM

The quality of structural detail and the ability to detect fluorescent reporters enabled by FDGN further allowed us to apply a quantitative approach to understand TEV-cell interactions by evaluating distribution of TEVs and TEV-associated fluorescent labels, FM1-43FX (blue) and PKH26 (red) fluorescent.

TEV distribution on the WI-38 cell surface at 2 or 24 hpa was represented by red dots on the respective SEM microgram (Fig. 4b*, TEV, left panel*). TEVs were widely distributed on the cell and often presented as clusters at 2 hpa, while becoming more sparsely distributed throughout the cell by 24 hpa (the density of TEV was 85.25 ± 89.90 /µm^2^ with 2 hpa and 7.37 ± 5.59 /µm^2^ at 24, p-value < 0.05) (Fig. 4b*, Density, left panel*).. The fluorescent signals in the cell at 2 hpa corresponded with sites of TEV clusters. By 24 hours post treatment, a limited number of TEVs were seen on the cell surface, however the fluorescent labels were broadly distributed within the cells, especially perinuclearly, indicative of TEV uptake. The area occupied by the fluorescent labels between 2 hpa and 24 hpa increased from 2.80 ± 2.22 % to 7.48 ± 4.25 % (p-value < 0.05) (Fig. 4b*, Area, right panel*). These analyses indicated that punctate fluorescent signals on the cell surface at 2 hpa dispersed broadly into the cells by 24 hpa, corresponding to early tethering to the cell surface followed by uptake and re-distribution within the cells.

### TEVs decrease microvilli membrane meshwork

To evaluate the capability of FDGN for cell surface imaging during the TEV initiated changes, we compared the topography of WI-38 cells treated with TEVs at 0, 2, or 24 hr post treatment via SEM. The cells at 0 hpa exhibited numerous microvilli and a randomly organized meshwork (Fig. 4c*, 0 hpa, left panel*). At 2 hpa, numerous TEVs appeared to be tethered to the cell surface and the meshwork revealed a modest rearrangement with a longitudinal orientation (Fig. 4c*, 2 hpa, 2^nd^ left panel*). By 24 hpa, only a limited number of TEVs were observed, whereas the meshwork showed a highly strict arrangement (Fig. 4c*, 24 hpa, 2^nd^ right panel*). Sporadic clusters of TEVs were readily found in depressions of the membrane. The density of microvilli was assessed at 0 hpa, 2 hpa, and 24 hpa. Compared to 0 hpa treatment, microvilli density decreased following TEV treatment, although there was no difference between 2 hpa and 24 hpa (0 hpa: 1.38 ± 0.16 /µm^2^; 2 hpa: 0.47 ± 0.08 /µm^2^; 24 hpa: 0.34 ± 0.18 /µm^2^; p-value < 0.05 for 0 hpa vs. 2 hpa as well as 0 hpa vs. 24 hpa) (Fig. 4c*,Microvilli density, right panel*).

### Effect of TEVs on WI-38 cell Meshwork

To examine the nature of the meshwork more closely, we used high power SEM and monochromatic inversion of high-power SEM images at the 0, 2, and 24 hr post-treatment time points. Control WI-38 cells (0 hpa) had numerous microvilli (*red arrow)*, and a relaxed meshwork of fibrous stipples (Fig. 4d*, 0 hpa*, *SEM, upper left panel*). Inverted SEM images revealed microvilli (*red arrow)* presented as dark rods, and each polygon created space between the fibers of the meshwork and was evenly open and relaxed without any indication of directionality or compression. The membrane pits (*blue arrow*) were not easily observed with SEM, but inverted SEM revealed the presence of bright and expanded space indicative of a membrane pit (Fig. 4d*, 0 hpa, Inverted SEM, lower left panel*). As expected, at 2 hpa, we observed numerous large (*large red arrowhead*) and small TEVs (*small red arrowhead*) on the cell surface with SEM (Fig. 4d*, 2 hpa*, *SEM*, *upper middle panel*). The TEVs appeared to be tethered to cell surface with thick fibers. The number of microvilli was limited but the number of membrane pits appeared to be increased (*blue arrow*). The inverted view highlighted a rearrangement of the fibrous network and membrane pits (*blue arrow*) compared to 0 hpa (Fig. 4d*, 2 hpa*, *Inverted SEM*, *lower middle panel*). The rearranged network was primarily associated with a longitudinal bi-directional extension and accommodated newly formed membrane pits with flexible expansions. Large (*large red arrowhead*) and small TEVs (*small red arrowhead*) and pits (*blue arrow*) were indicated by dark spheres and bright open space in the inverted view, respectively. TEVs were generally found in the proximity of membrane pits.

By 24 hpa, the cells displayed sparsely distributed small and large TEVs on the surface (Fig. 4d*, 24 hpa, SEM, upper right panel*). Here again, large (*large red arrowhead*) and small TEVs (*small red arrowhead*) were prominently featured in the proximity of membrane pits, which appeared as either a cup shape or depression (*blue arrow*). At this time point, the fibrous meshwork had undergone significant rearrangement with prominent fibers (>25nm) displaying a strict longitudinal orientation and associated fine fibers (<20nm) running across. This highly ordered organization was easily observed by SEM or inverted SEM. Meanwhile the relationship among active elements of the cell surface, large (*large red arrowhead*) and small TEVs (*small red arrowhead*), and membrane pits (*blue arrow*) was easily appreciated by the inverted SEM, where the meshwork was altered due to the presence of large membrane pits and TEVs were circumferential or adjacent to the pits. The overall fibers of meshwork were strictly oriented in a longitudinal direction (Fig. 4d*, 24 hpa*, *Inverted SEM, lower right panel*).

### TEV-host cell interaction

The capability of FDGN to preserve detailed structures and retain fluorescent signals allowed us to apply SEM CLEM to further study the direct interactions between TEVs and host cells and its sequelae. WI-38 cells at 2 hpa exhibited both the FM1-43FX (blue) and PKH26 (red) fluorescent probes that were used to label TEVs (Fig. 5a*, FLM, upper left panel*). SEM examination of the identical field revealed a large and dense speckled cluster on the cell surface (Fig. 5a*, SEM, upper 2^nd^ left panel*). The merged image of FLM and SEM demonstrated that the fluorescent signal co-localized with the speckled clusters (Fig. 5a*, Merged, upper 2^nd^ right panel*). A closeup of a selected area (*inset*) further demonstrated that the fluorescent labels were closely associated with speckles, confirming these were TEVs (Fig. 5a*, Up-close, upper right panel*, *double arrowheads*). At 24 hpa, FLM revealed widely distributed fluorescent signals (Fig. 5a*, FLM, middle left panel*). SEM of this field showed that the cell surface (Fig. 5a*, SEM, middle 2^nd^ left panel*) was mostly devoid of speckles. A merged image of FLM and SEM indicated that both fluorescent signals were broadly distributed within the cell (Fig. 5a*, Merged, middle 2^nd^ right panel*). There was a patchy distribution of fluorescence in a region of the cytoplasmic extension (*inset*). The magnified view of the inset revealed that this patchy fluorescence closely overlapped with the membrane pits on the cell membrane (Fig. 5a*, Up-close, middle right panel*, *blue arrow*).

**Fig. 5.**
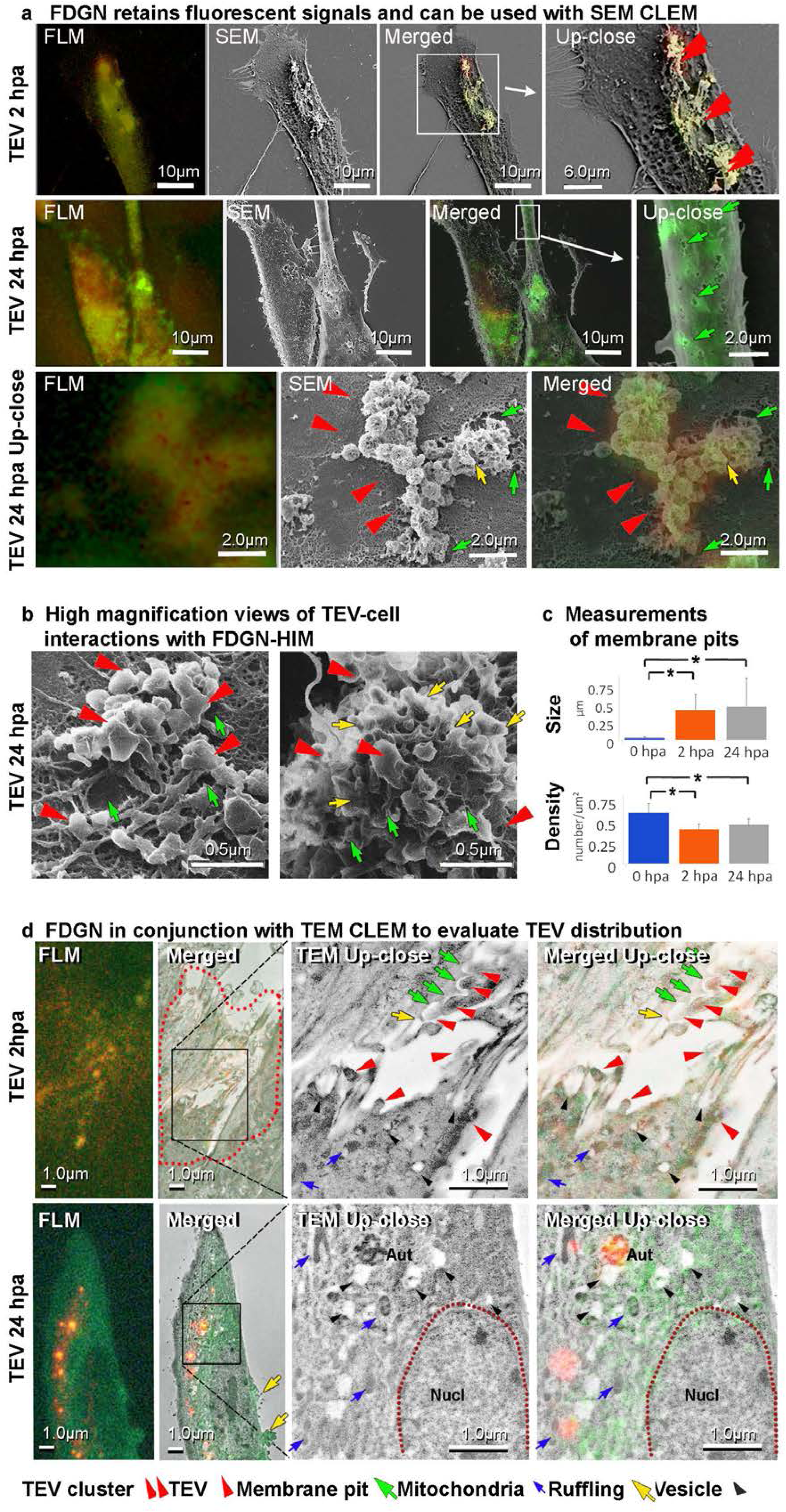
FDGN allows visualization of TEV-WI-38 cell interactions. **a,** FDGN processing retains fluorescent signals and can be used with SEM CLEM An **FLM** micrograph at **2 hpa** shows a cluster of well retained FM1-43FX (green) and PKH26(red) positive signals (***top left panel***). **SEM** micrograph of the ROI shows accumulated small spherical objects in an identical area (***top 2^nd^ left panel***). A **Merged** view revealed FM1-43FX and PKH26 positive signals present on the cell in the SEM micrograph (***top 2^nd^ right panel***, ***inset***). Higher magnification of the area in the inset reveals that positive fluorescent signals represent clusters of TEVs (***top right panel, double arrowheads*** 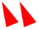). At 24 hpa **FLM** micrograph (***middle left panel***) shows widely distributed fluorescent signals. **SEM** microgram of the ROI reveals that the cell surface is mostly devoid of small spherical objects (***middle sencond left panel***). The **Merged** view shows the broad distribution of fluorescent signals within these cells (***middle 2^nd^ right panel***) and a peripheral part of the cell (***inset***) shows patchy fluorescence. A higher magnification of the **inset** reveals the association between the fluorescence and membrane pits (***blue arrow*** 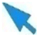)(***middle right panel***). The **FLM** at **24 hpa Up-close** (***lower left panel***) shows a cluster of well-retained FM1-43FX and PKH26 signals. This ROI view by **SEM** reveals a cluster of globular objects (***red arrowhead*** 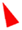) and membrane ruffling (***yellow arrow*** 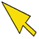) tethered on the cell surface (***lower middle panel***) with adjacent membrane pits (***blue arrow*** 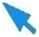). The **Merged** image reveals that these globular structures (***red arrowhead*** 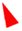) and membrane ruffling (***yellow arrow*** 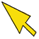) contain TEV-associated fluorescent markers (***lower right panel***). **b,** Detailed views of TEV-cell interactions with FDGN-HIM WI-38 cells underwent FDGN treatment prior to HIM imaging. HIM image shows a up-close view of the juxtaposed TEVs (***arrowhead*** 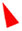) and adjacent membrane pits (***left panel***). Another image shows an area with a high concentration of fusion of TEVs along with host cell membrane ruffling (***yellow arrow*** 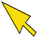) near the site of TEV-cell interaction and adjacent membrane pits (***blue arrow*** 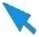)(***middle panel***). **c.** TEV impact on cellular membrane pits The changes in the **Size** of diameter (µm) and the **Density** of membrane pits (per µm^2^) between WI-38 cells at **2 hpa** and **24 hpa** were quantified. Upper panels show the size of membrane pits in cells with control (***0 hpa***) and **2 hpa** and **24 hpa**. Lower panels shows changes in the density of membrane pits in each condition. * Indicates statistical significance, with p-value < 0.05. **d, FDGN** in conjunction with TEM-CLEM to evaluate TEV distribution WI-38 cells treated with TEVs underwent FDGN and were further processed for TEM CLEM. Ultrathin sections of cells at **2 hpa** retain both fluorescent signals (**2 hpa, *FLM, upper panel***). The ROI of the section was imaged using TEM and the merged image (**2 hpa, *Merged, upper panel***), reveals well-preserved ultrastructure and fluorescent signals on the cell surface and submembrane region (***dotted red line***). Higher magnification image of the selected area (***inset***) shows the presence of TEVs attached on the cell surface ***(large red arrowhead*** 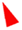), some of which are adjacent to membrane pits (***blue arrow*** 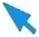) or membrane ruffles (***yellow arrow*** 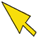), as well as vesicles ***(black arrowhead*** 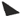) and mitochondria (***blue arrow*** 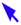) in the cytoplasm close to the surface of the cell (***TEM Up-close, upper panel***). The FLM/TEM **Merged** close-up image shows that fluorescent signals co-localize with TEVs, vesicles, and mitochondria (***Merged Up-close, upper panel***). Similarly, FLM microgram of the ultrathin section from the samples at **24 hpa** shows fluorescent signals on the TEVs attached to the cell surface, a large part of the cytoplasm (**24 hpa**, ***FLM, lower panel***). In combination with TEM, the fluorescence signal was observed on the cell surface TEV, cytoplasm, and small area of the nucleus (***Merged, lower panel***). Higher magnification TEM image of the selected area (***inset***) reveals the presence of vesicles, mitochondria, and autophagosome in the proximity of the nucleus (***TEM Up-close, lower panel***). Merged images of TEM/FLM (***Merged up-close, lower panel***) reveals that fluorescent signals co-localize with mitochondria, vesicles, autophagosome, as well as peripheral area of the nucleus.

We used higher magnification to examine the TEVs on the cell surface at 24 hpa. The FLM showed a cluster of signals of both FM1-43FX (blue) and PKH26 (red) (Fig. 5a*, FLM, lower left panel*). The SEM view of the specific field revealed a cluster of spheres (*large arrowhead*) and an aggregate of membrane ruffling (*yellow arrow*) (Fig. 5a*, SEM, lower middle panel*). As indicated, membrane pits (*blue arrow*) were in the proximity of the clustered spheres. The merged view confirmed that both spheres (*arrowhead)* and membrane ruffling (*yellow arrow*) were associated with fluorescent signals (Fig. 5a*, Merged, lower right panel*). This observation indicated that the clusters of TEVs interacted with the cell via membrane pits or ruffling. Thus we confirmed that the spheres tethered to WI-38 cells expressed the TEV fluorescent markers and TEV-host cell interactions were associated both with the formation of membrane pits and ruffling and together suggest multiple modes by which TEVs interact with host cells.

### FDGN/ HIM analysis of TEV cellular interactions

Cells prepared by FDGN at 24 hpa were subjected to super-resolution scanning HIM imaging without a conductive coating. FDGN revealed notably distinct profiles of membrane pits (*blue arrow*), TEVs (*arrowhead*), and meshwork on the membrane. At times, a cluster of TEVs were juxtaposed with a number of large membrane pits (Fig. 5b*, left panel*). At other times TEVs fused with the ruffled cell membranes (Fig. 5b*, right panel*, yellow *arrow*). In this case, small membrane pits appeared at fusion sites (*blue arrow*) and together suggests multiple methods of TEV entry.

### Quantitative analyses of membrane pits

We employed FDGN to quantitatively evaluate the dynamic nature of the cellular topography. First, we analyzed changes in membrane pits at 0 hpa, 2 hpa and 24 hpa (Fig. 5c). We observed an increased diameter of membrane pits with TEV application compared to control but no further increases between 2 hpa and 24 hpa (*upper panel, Size*, 0 hpa: 0.028 ± 0.013 µm, 2 hpa: 0.042 ± 0.21 µm, 24 hpa: 0.47 ± 0.40 µm, p-value < 0.05 for 0 hpa vs. 2 hpa and 0 hpa vs. 24 hpa). We also observed a decreased density of membrane pits following TEV application and control (0 hpa) but no further differences between 2 and 24 hpa (Fig. 5c*, right lower panel, Density,* 0 hpa: 0.62 ± 0.11 /µm^2^, 2 hpa: 0.42 ± 0.06 /µm^2^, 24 hpa: 0.47 ± 0.08 /µm^2^, p-value < 0.05, 0 hpa vs. 2 hpa and 0 hpa vs. 24 hpa). Furthermore, we noticed a larger variability in the size of membrane pits with TEV applications as the treatment time increased (0 hpa: < 0.066 µm, 2 hpa: <1.4 µm, and 24 hpa: <3.1 µm.) suggesting dynamic TEV-cell interactions give rise to membrane pits. These quantitative analyses of microvilli and membrane pits illustrate the distinct reorganization of cellular topography following interactions of TEVs with WI-38 cells.

### Application of FDGN for TEM CLEM

Finally, we tested the applicability of FDGN for TEM CLEM in WI-38 cells, at 2 hpa and 4 hpa. Cells initially dehydrated for SEM with FDGN and further processed for embedding in plastic resin and ultrathin sections (90nm) were analyzed by FLM and followed by TEM.

At 2 hpa, cells exhibited a punctate pattern of fluorescent signals from both the TEV FM1-43FX and PKH26 probes (Fig. 5d, *2 hpa, FLM*). Further examination of the region of the section revealed well-preserved ultrastructures, while the TEM merged image indicated that the fluorescent signals were primarily located at the cell surface and within the cytoplasm adjacent to the cell surface (Fig. 5d, *2 hpa, Merged*, *outlined by the dotted red line*). The ultrastructure of the selected area (*inset*) shown in the TEM view, revealed prominent vesicles associated with the external surface (*large red arrowhead*), some of which are facing membrane pits (*blue arrow*) or ruffling (*yellow arrow*), and the submembrane area (*black arrowhead*) (diameter of vesicles 193.13 µm ± 28.48) (Fig. 5d, *2 hpa, TEM Up-Close*). Mitochondria (*blue arrow*) were also observed. A high magnification view of the merged image revealed the fluorescent signals associated with TEVs were on the cell surface (*red arrowhead*) and on vesicles within the cytoplasm (*black arrowhead*) as well as mitochondria (*blue arrow*) (Fig. 5d, *2 hpa, Merged Up-close*). Membrane pits (*blue arrow*) or ruffling (*yellow arrow*) were noted in the positions adjoining vesicles on the cell surface.

At 24 hpa a broad distribution of both fluorescent signals was visible via FLM (Fig. 5d, *24 hpa, FLM*). The merged FLM and TEM image (Fig. 5d, *24 hpa, Merged*) displayed both fluorescent signals on clusters of TEVs on the cell surface with a ruffled membrane (*yellow arrow*) and a more broad distribution within the cell body, but prominent in the perinuclear region. This image also showed that the fluorescent signals were expressed in limited parts of the nucleoplasm and the ultrastructure of the selected area (*inset*) is highlighted (Fig. 5d, *24 hpa, TEM Up-close*). There were noticeable vesicles, with a diameter of 217.22 µm ± 33.67, in the same size range with those measured at 2 hpa. A merged close-up showed vesicles, mitochondria, and autophagosomes and a limited part of the nucleoplasm associated with fluorescent signals (Fig. 5d, *24 hpa, Merged Up-close*, *arrowhead, Mit, Aut*).

Analyses of TEM CLEM between 2 hpa and 24 hpa revealed that the FM1-43FX (blue) and PKH26 (red) TEV signals were actively redistributed from the cell surface to the cytoplasm as well as to a portion of the nucleus. In addition, some of the TEV fluorescent markers on the cell surface colocalized with membrane pits and ruffling. These observations were complementary to those found using SEM CLEM. Moreover at 24 hr after treatment, both fluorescent signals were not limited to TEVs and variably associated with other membrane structures in the cytoplasm, suggesting the possibility of direct TEV association with these organelles. This observation suggests that TEVs maintain their vesicular form after entry into the cell and interact with intracellular structures. However, it is also important to note that TEVs are likely degraded over time and that the membrane-associated fluorescent dyes may be re-cycled and potentially label other intracellular compartments.

## Discussion

FDGN has produced promising resuts wiht unique and intricate cytoarchitecture of the cytoplasmic membrane and retained clear fluorescent signals as we reported earlier (Dann et al., 2018; Santulli et al., 2014). The combination of high-quality structural data and fluorescent reporter signals provides a powerful new method for high-resolution SEM CLEM, enabling the detection of membrane structures and interactions that previously were not appreciated. Furthermore, we showed that FDGN can be applied to HIM, and TEM CLEM, thereby allowing detailed analyses by multiple imaging systems.

The present study confirmed that the ethanol dehyderation compromised the surface cell membrane integrity and OsO_4_ mitigated (Saunders et al., 1968). Intriguingly, we found that cytoarchitecture could be further improved by omitting these two chemicals altogether and subjecting samples to FDGN processing. To avoid processing artifacts, preparing samples without ethanol or OsO_4_ treatment and viewing the samples in a hydrated state is considered an ideal approach (Ogura, 2019; Takaku et al., 2020). A few studies have reported such imaging methods for SEM CLEM (Thiberge et al., 2004; Wojcik et al., 2015). These SEM CLEM methods visualized fine cytoskeleton and organelles and preserved the distribution of fluorescent reporter molecules; however, interactions between the cell surface and its external environment in detail have not been shown. FDGN provided an alternative solution by preserving molecular organization and cytoarchitecture by minimizing chemical interventions and uniquely allowing longitudinal imaging of the mechanisim of interactions of HCV virons or TEVs with the cell membrane at specific time points. Furthermore, it differnciated cell type-specific morphological properties in between WI-38 fibroblasts and Huh7.5 hepatocytes.

By correlating with subcellular distribution of fluorescent reporters, we reliably imaged the accompanying dynamic structural changes, nuclei, including bulged nuclei with RFP in HCV-infected Huh7.5 cells or with TEV interactions with WI-38 cells. The present study connected our understning of the effect of HCV-host cell interaction with drastic deformation of nuclear organization reported (Allen et al., 2008; Cortese et al., 2017; Cotter et al., 2007; Denais et al., 2016; Kamikawa et al., 2022; Kiseleva et al., 2001; Lee et al., 2012; Lindenboim et al., 2020; Robijns et al., 2018). By doing so, it also allowed for recognition of other associated structural changes, including reorganization of the cytoskeleton, microvilli, and membrane pits, which are considered equivalent to the specialized coated membrane pits, involved in receptor-mediated endocytosis. Together FDGN with SEM CLEM and TEM CLEM enhanced our understanding of how HCV virions and TEVs interact with and enter respective host cells, and glimpsed their subsequent behavior within the cell.

We believe that SEM could expand its important role as it allows direct imaging of detailed interactions at the cell surface and FDGN processing presented the images closely reflected our current understanding of pathogen or TEV interaction with its host cell and cell entry (Blanchard et al., 2006; Colpitts et al., 2020; Lindenbach and Rice, 2013). From technical point of view, each step of FDGN work flow, including glutaraldehyde fixation in sodium cacodylate buffer, washing in water, plunge freezing and CPD, is reasonable and accepted in the practice in the field of EM and the additional applicataion of nitrogen gas flow is another gentle step. We expect that continuous studies will validate the role of FDGN.

Based on the present study, our understanding of the HCV- and TEV-host cell interactions are summarized in the schematic diagram (Fig. 6a, HCV, left side of panel). A mock-infected human hepatoma cell displays a flat cell profile with a relaxed cytoskeletal meshwork (Mock). HCV virions initiate interaction and tether to the cell membrane. HCV particles interact with cognate receptors, triggering reorganization of the cytoskeleton, and the formation of membrane pits and more microvilli in the cell, followed by HCV entry into the cell by clathrin-mediated endocytosis (HCV entry). Infection triggers a dramatic rearrangement of the nucleoplasm and generation of a nuclear bulge (Post entry). HCV particles enter the cell via clathrin-mediated endocytosis and replicate in the endoplasmic reticulum (ER). TEVs undergo a similar process with some variations (Fig. 6a, TEV, right side of panel); e.g., whereas HCV increased microvilli in Huh7.5 cells, TEVs decreased microvilli in WI-38 cells, suggesting particle-specific interactions. A relaxed cytoskeleton of the host cell (0 hpa) undergoes rearrangement upon TEV interaction, in a receptor-dependent or independent manner. The cell allows TEVs to tether to the cell surface (TEV tethering). TEVs subsequently enter host cells possibly through proximal membrane pits (via clathrin-dependent endocytosis) or membrane ruffles (Ruf) (via clathrin-independent macropinocytosis) (TEV entry). The fibroblast subsequently undergoes dynamic morphological transformation with formation of a tightly organized bi-directional cytoskeletal network, resulting in an elongated cell appearance. Meanwhile, TEVs interact with endosomes and release their cargo. They can also be transferred to lysosomes/autophagosomes directly or indirectly. After TEV entry, the cell may maintain membrane pits with TEV-associated fluorescent signal for periods of time and their lipid-rich membranes can be recycled and re-used by the cell (post entry). Using FDGN, we observed subtle differences between TEV-host cells interactions through both membrane pits or membrane ruffling. Determining whether small sized TEVs enter the cell either through endocytosis, or macropinocytosis, would require more sensitive and specific markers. Nonetheless, our observations are in agreement with the understanding that HCV enters via clathrin-mediated endocytosis (Blanchard et al., 2006; Colpitts et al., 2020; Lindenbach and Rice, 2013) and TEV’s can employ a variety of methods, including non-specific macropinocytosis or via specific, receptor-dependent pathways, including endocytosis by either clathrin-dependent or clathrin-independent routes (Mulcahy et al., 2014).

**Fig. 6.**
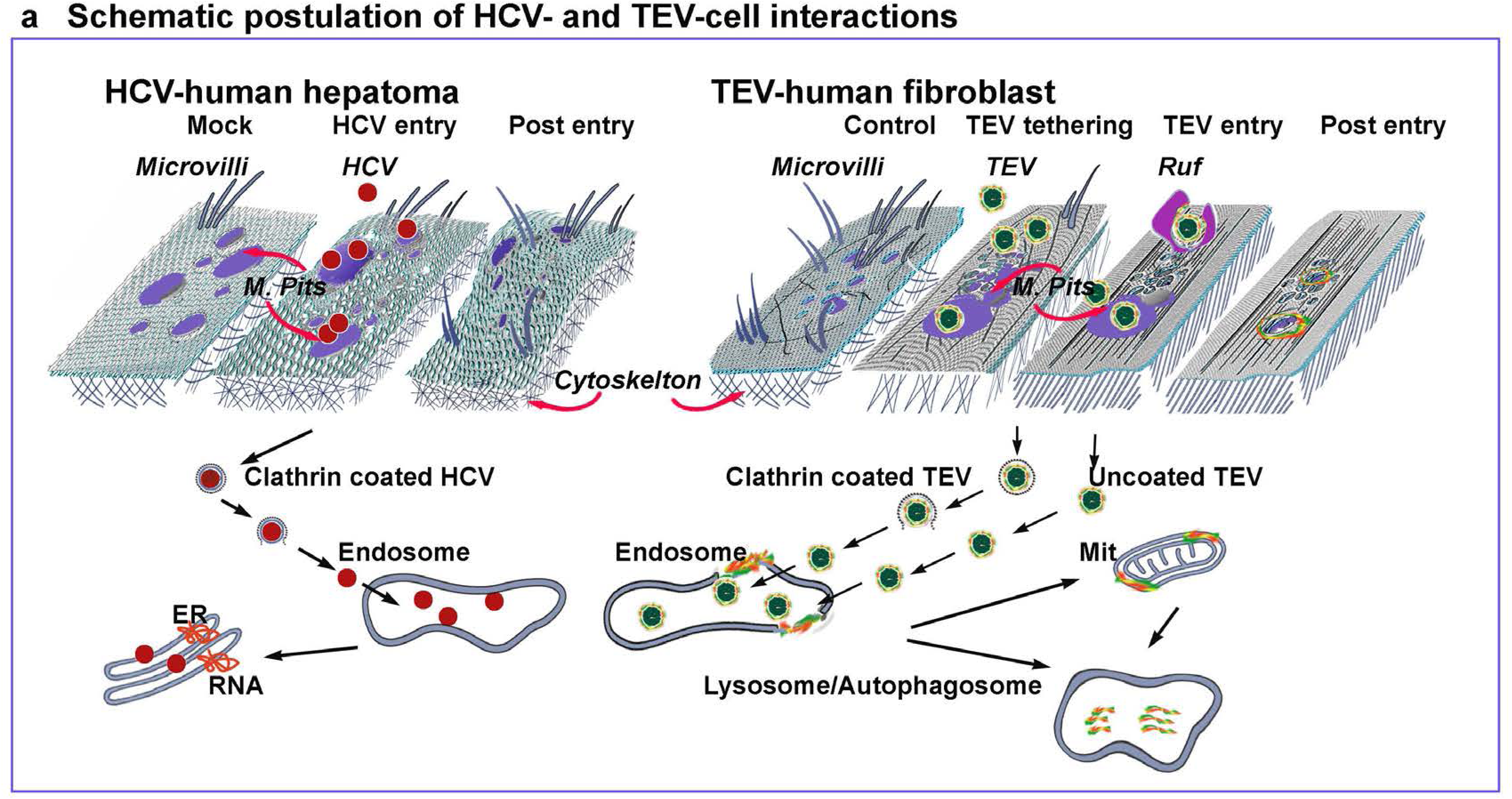
a, Schematic diagram of viral and TEV-cell interactions. The right panel shows our understaning of how HCV-Huh7.5 cell interact with (***HCV***). The mock infected cells display a relaxed cytoskeleton and as the HCV virion contacts the cell, receptor interactions initiate actin-driven membrane rearrangements, which in turn leads the HCV virion to tether to the cell surrounded by a depressed membrane with subsequent entry through a membrane pit (***M. Pits***), in a clathrin-mediated endocytic manner (***HCV entry***). After viral entry, the cell rearranges the nucleoplasm (***Post entry***). Meanwhile, through the engagement with the endosome, the virus releases its RNA to the ER to initiate its reproduction. The right panel shows our perspective of how cell interacts with TEVs. Cells not exposed to TEVs display many microvilli and a relaxed cytoskeleton meshwork. TEV-cell interactions can be initiated via integrin-mediated contact, which facilitates TEV tethering (***TEV tethering***). In fibroblasts the tethering of TEVs initiates actin-dependent changes in membrane organization and cell shape. This results in either membrane pit (***M. Pits***) formation or ruffling (***Ruf***) to support TEV entry (***TEV entry***). After the TEV entry, membrane pits and TEV-associated fluorescent dye can be seen at the entry sites (***Post entry***). TEV entry can occur via clathrin-dependent endocytosis or a clathrin-independent macropinocytosis. In each case subsequent interactions with the endosome allow TEVs to release its cargo. Cytoskeleton arrangement transforms fibroblasts into a directionally organized elongated cell.

A recent report described how microvilli and the cytoskeleton coordinately respond to the cell’s external environment. For example, microvilli serve as a signaling center for receptor-based interaction in T cells (Mastrogiovanni et al., 2020; Mathieu et al., 2019). Microvilli are enriched in the presence of T-cell receptors (TCR) and co-receptors, and could serve as a compartment to reorganize TCR signaling related proteins (Mathieu et al., 2019) and subsequently drive actin cytoskeleton remodeling for immunological synapse formation (Mastrogiovanni et al., 2020). It is plausible that microvilli of hepatocytes and fibroblasts contain distinct receptors to respond to specific pathogens. Furthermore, cell membrane fusion and membrane pit formation are closely linked to actin activity in experimental membrane fusion models (Cogger et al., 2013; McGuire et al., 1992). Our study indicated HCV interaction stimulated Huh7.5 cell microvilli formation while TEV reduced this process in fibroblasts. Whether HCV-induced microvilli enhances the cell’s ability to interact with the immune system, while reduced microvilli following TEV interactions leads to reduced interactions with immune cells is not known. Recent work on SARS-CoV-2 replication in human bronchial epithelium revealed a transient and rapid loss of the ciliary layer and structural flattening of microvilli, which might be explained by axoneme loss and misorientation of remaining basal bodies (Caldas et al 2020). Thus, these alterations were thought to be indirect products of the interaction (Robinot et al., 2021). Our observations in Huh7.5 cell and WI-38 cells indicated that the initial rearrangement of microvilli, meshwork and membrane pits and overall modification of cell shape arose through interactions between HCV or TEVs prior to entry and likely facilitates entry into cells. The mechanisms of pathogen-host cell interactions however are not a singular process and understanding the precise entry processes used by viruses or TEVs will be the subject of future studies.

Existing studies have suggested that TEVs can release their contents into the cytoplasm through an unidentified mechanism (Bonsergent et al., 2021; Joshi et al., 2020), but the fate of the cargo is not known and is a focus of ongoing studies. We previously showed using immuno EM that TEVs carrying mitochondrial DNA were sequestered and fused with mitochondria suggesting that TEVs may release their cargo at specific locations within recipient cells (Sansone et al., 2017). Our TEM CLEM analysis in the present study revealed TEV-related fluorescent dye in the mitochondria, supporting the notion of direct TEV-mitochondrial interactions. Sequestered TEVs appear to be retained in endosomal vesicles for a certain period before releasing their contents within the cell. However, due to the nature of lipid affinity dyes, recycled membranes could also result in a re-distribution of fluorescent-labeled membranes throughout the cell or accumulation in lysosome/autophagosomes. Nonetheless, there is increasing evidence that TEVs are involved in a broad range of physiological and pathological processes, including cellular homoeostasis, cancer metastasis, and cardiovascular diseases (Busatto et al., 2021; Rigalli et al., 2020). Given TEVs’ ability to travel systemically in plasma and be taken up by a variety of distal cell types, recent studies have suggested that TEVs can be modified to act as superior synthetic drug delivery vectors (Herrmann et al., 2021). The improved imaging made possible by the use of FDGN processing in combination with advanced microscopy will be a valuable tool for evaluating the fate of endogenous TEVs and those modified for drug delivery.

Indeed, the detailed examination of cytoarchitecture in conjunction with fluorescent reporters with SEM CLEM is novel and provides a previously unattainable insight into events occurring on cell surfaces as well as within intracellular organelles. Although fluorescent microscopy can reveal time-sensitive localization of molecules, it does so at the expense of higher resolution. Moreover, while TEM provides a higher resolution of organization of the sub-membranous cytoskeleton, it is limited to thin sections that preclude the ability to visualize larger cellular areas and interactions with virions or TEVs. To extend TEM data to 3D reconstruction, more highly specialized instruments and time are required. Thus, SEM CLEM with FDGN provides a high resolution, user-friendly approach to study remodeling events on the cell membrane induced by cellular interactions with pathogens or TEVs. This approach may be highly applicable to many viral infections including SARS-CoV-2 (Caldas et al., 2020), as it would allow detailed examination of interactions of the virus with cells with a preserved, dynamic structural cellular architecture that in turn, could inform our understanding of cellular responses to virus invasion. Ultimately, this information could be applied to develop novel therapeutic approaches to limit viral infection and/or replication (Duncan et al., 2020). In conclusion, we demonstrate that FDGN processing can be applied to biological samples under physiological and/or pathological conditions and can provide invaluable information for understanding dynamic cellular interactions with TEVs or viral pathogens.

## Score system

+++: excellent quality

++: intermediate quality

+: poor quality

-: not detectable

n/a: not tested

## Materials & Methods

### Mammalian cell culture

Human hepatoma derived Huh-7.5 cells expressing a RFP-NLS-IPS construct(Jones et al., 2010) were cultured at 37°C, with 5% CO_2_ in Dulbecco’s Modified Eagle Medium (DMEM, Invitrogen, Carlsbad, CA) containing 10% fetal bovine serum (FBS) and 0.1 mM nonessential amino acids for 24 hours. Cells were grown on a pair of plastic coverslips (ACLAR 33C film, Electron microscopy sciences, Hatfield, PA) gridded by carbon film and subjected to HCV RNA–electroporation or vector (mock) treatment and fixed at 24 hpi(Jones et al., 2010). Fluorescent nuclear stain (NucBlu R37605, Thermo Fisher Scientific, Waltham, MA, USA) was added to identify the location of nuclei. Human lung fibroblasts, WI-38 (ATCC), were maintained in alpha-modification Minimum Essential Medium, supplemented with 10% exosome-depleted fetal bovine serum (FBS) (Gibco, Thermo Fisher Scientific, Waltham, MA, USA) and penicillin-streptomycin. We incubated WI-38 cells with exosomes isolated from 4175 lung tropic MDA-MB231 human breast cancer cell cultures on ACLAR coverslips gridded by carbon film and labeled exosomes with the lipophilic dyes FM1-43FX (green) (ThermoFisher, Waltham, MA) or PKH26 (red) (Sigma-Aldrich, St. Louis, MO)(Hoshino et al., 2015). Cells were fixed at 0 (without TEV application), 2 or 24 hpi, and used for various studies described below.

### Microscopy

All samples were initially rinsed in the respective serum free culture medium and fixed in 0.075M sodium cacodylate buffer (pH 7.4), containing 2% paraformaldehyde, 2.5% glutaraldehyde, 0.2M sucrose, 5mM magnesium chloride, and 2mM calcium chloride. Cells were examined under a fluorescent microscope to ensure that the samples were free of contamination and then washed in sodium cacodylate buffer (5min, 3 times) and water (5min, 3 times). One of the following four procedures were subsequently conducted: CSEM: samples were post-fixed in ice-cold 1% OsO_4_ in 0.075M sodium cacodylate buffer for 1 hour, treated with increasing concentrations of ice-cold ethanol (15 min each in 50%, 75%, 95%, and 3 times in 100%) and followed by the conventional CPD procedure, initiated by filling the CPD chamber with 100% ethanol (Autosamdri A-815, Series A, Tousimis Research) and running the automated program according to the manufacturer’s instructions. 2) FDGN: this procedure is initiated by a full absorption of liquid from the coverslips (cut into an approximately 3 x 3mm sized ACLAR sheet) with filter paper, followed by freezing in a liquid ethane cup pre-cooled in a LN_2_ bath. (Fig. 1) Coverslips were then transferred to a sample holder located in a pre-colled GN_2_ filled chamber, with a 10cm petri dish filled with silica gel (Fisher Scientific, Cat# S679-500) covered by lint-free linen (Ted Pella, Cat# 812-14), dehydrated with a continuous flow of GN_2_ (at - 140 °C) (AFS2 from Leica Microsystems Inc., Vienna, Austria) overnight and gradually increased to −30 °C. It was subjected to an ethanol-free CPD procedure filled with GN_2_, a modification of existing method (Joens et al., 2013). It is critical to avoid condensation throughout the procedure, in particular during the transfer from the GN_2_ chamber to the CPD unit, thus, we devised a custom-made cryogenic transporter and GN_2_ applicator (Fig. 1). A cryogenic sample transporter was prepared by a form box. Inside the box, a SEM sample holder is placed on a metal platform in a glass beaker which is secured by two horizontal arms made of foam. The sample holder is protected by a plastic safty cup and placed on the metal platform. To ensure proper sample preservation, the bottom half of the unit is filled with LN_2_ and the upper half is filled with GN_2_ and the sample is kept cold to avoid condensation. The GN_2_ applicator set up is prepared with a Pyrex glass flask and a connected silicon tube (inner diameter: 6.4mm; outer diameter; 9.6mm) through a rubber top. The flask is half-filled with LN_2_ and collect a continuous flow of cold GN_2_ by gently placing the rubber top that conneted to a silicon tube not to cause an excessive pressure. While Pyrex glass is tolerate extreme temperature between −192 and +500 °C and consider suitable to use of LN_2_, the flask was placed behind a shield for an unexpected incidents. When GN_2_ flow wanes, topple additional LN_2_. The samples in the GN_2_ chamber was translocated to the cryogenic transporter with precooled plastic safety cup and outfitted with a lid. For the sample transfer to the CPD chamber, the chamber (3.18 cm in diameter) was pre-cooled according to the manufacturer’s instructions and by the addition of cold GN_2_ from the GN_2_ applicator for a minimum of 5 min to reach to −30 °C. The lid of CPD chamber should be placed on top of the silicon tube of the GN_2_ applicator to maximize cooling efficiency. For the use of the CPD at sub-freezing temeperture, Acetone Package Kit with the suitable teflon O-ring and spanner knurl nuts (Tousimis, Rockville, MD, #8770-91) was used. The cryogenic transporter was located in proximity to the *CPD chamber* and the sample holder was subsequently transferred into the CPD chamber. It was critical to maintain a continuous application of cold GN_2_ to the sample holder by keeping the end of the silicon tube directly over the sample holder during this transfer. Upon relocating the sample holder, the O-ring and lid were replaced to proceed with the closure. During this time, the silicon tube remained inserted into the chamber to continuously supply cold GN_2_ and removed only at the final stage of tightening screws for the lid. Once closed the CPD program was initiated by introducing liquid CO_2_. 3) CSEM-Os: The samples were treated in the same way as CSEM method, except OsO_4_ post-fixation was omitted. 4) CPD-ctr: The cells underwent the same procedure as in FDGN, but the CPD steps were omitted and instead, the GN_2_ chamber temperature was slowly raised to +23 °C (at a rate of ∼10 °C/ hour) at the end of overnight incubation. After the dehydration step, samples were subjected to LM (Eclipse with DS-Fi1 digital imaging system with NIS-Elements F 4.6, Tokyo, Japan), SEM (Leo 1550, 10eKV, 10µm aperture, ∼100pA, Smart SEM v 5, with 5nm thick iridium coating, Carl Zeiss, Oberkochen, Germany), or HIM (Orion, ∼30eKV, Smart SEM v 5, 5µm aperture, ∼0.5pA) at Carl Zeiss, Peabody, MA, examination. For HIM, uncoated samples were used in the presence of an electron flood gun which allows for imaging without a conductive coating and provides superior detail.

For TEM examination, samples were incubated in a freeze-substitute, containing 0.2% uranyl acetate and 95% of acetone, for 4 hours and followed by Lowicryl HM20 for 24 hours with agitation and polymerized under UV exposure at −45 °C in an automated processor (AFS2, Leica Microsystem Inc., Vienna, Austria). Ultrathin sections (90 nm thickness) were examined with a Nikon Eclipse and JEOL 100CX TEM at 80KV with a digital imaging system (XR41-C, Advantage Microscopy Technology Corp, Denver, MA). All sample preparations and LM and EM data collection were conducted at the Electron Microscopy Resource Center at the Rockefeller University, New York, NY. HIM data collection was conducted at the Ion Microscopy Innovation Center, Zeiss Microscopy LLC, Peabody, MA.

### Quantitation and statistical analyses

For quantitative analysis, using the RFP-NLS-IPS for HCV infection or lipophilic dyes FM1-43FX (green) (ThermoFisher, Waltham, MA) or PKH26 for EV, representative cells were selected from images obtained under each condition. Microvilli on hepatoma cells and TEVs on fibroblasts were counted using Adobe Photoshop (2020, 21.1.0). Membrane pits of hepatocytes (Huh-7.5) (n=3) and fibroblasts (WI-38) (n=3), as well as fluorescent areas, were analyzed using ImageJ (1.52a) (NIH/LOCI, University of Wisconsin, Madison, WI). We used a two-tailed, unpaired Student’s t-test for two-group comparisons and one-way or two-way ANOVAs and post-hoc Bonferroni Correction for multiple-group comparisons. In all experiments, p < 0.05 indicated a statistically significant result. Bar graphs depicted mean values and the error bars in bar plots represented standard deviations of the mean (±SDM).

## Other abbreviations

FLM: fluorescence light microscopy
CPD: critical point dry
HIM: helium ion microscopy
SEM: scanning electron microscopy
TEM: transmission electron microscopy

## Acknowledgments

The authors thank Mark Ebrahim and Kris Tuthill for technical support and discussions and the members of Lyden Lab for valuable support. The current study was supported by The Rockefeller University, NIH R01 AI072613 (Rice), NIH R01 CA057973 (Rice). The authors gratefully acknowledge support from the Thompson Family Foundation (to D.L.), the Pediatric Oncology Experimental Therapeutics Investigator’s Consortium, the Malcolm Hewitt Weiner Foundation, the Manning Foundation, the Sohn Foundation, the AHEPA Vth District Cancer Research Foundation, the Children’s Cancer and Blood Foundation (to D.L.)

## Authors’ contributions

K.U. designed the methodology, carried out the test procedures and conceptualized the study, N.S., T.S., M.T.C., A.Ho, and A.Ha design and carried out experiments; C.H. performed data collection; K.U. made the draft of manuscript; E.M., C.K.,N.B and J.P. reviewed the data and manuscript; K.U., D.L., and C.R., offered financial contributions and provided overall supervision.

